# Coronavirus circulation and cross-species transmission in European bats: a new paradigm in betacoronaviruses

**DOI:** 10.64898/2026.10.13.758892

**Authors:** Jordi Serra-Cobo, Abir Monastiri, Carlos Herrera-Ubeda, Nuri Llorens Bàrbara, Maria I Costafreda, Albert Carcereny, Marc López-Roig, Jordi Garcia-Fernandez

**Author notes:** Correspondance: Jordi Serra-Cobo & Marc López-Roig.

## Abstract

Bats are major reservoir hosts of coronaviruses (CoVs), yet the ecological mechanisms governing CoV circulation and cross-species transmission remain poorly understood, particularly in Europe. Here, we investigated the diversity, host associations, and interspecific transmission dynamics of alpha- and betacoronaviruses in European bats using an integrated phylogenetic and network-based approach. We analysed 835 guano samples collected from 31 bat roosts in Catalonia and the Balearic Islands (Spain) between 2020 and 2024, encompassing 14 bat species from three families. CoV RNA was detected in 18.9% of samples, with five alphacoronavirus and two betacoronavirus subgenera identified. Virus–host networks were highly structured and non-random, with strong modularity and specialization, and a small subset of bat species occupying central connector roles. Notably, we report the first confirmed detection of sarbecoviruses in *Miniopterus schreibersii*, expanding their known host range beyond Rhinolophidae, and identify merbecoviruses in *Rhinolophus euryale* for the first time. Phylogenetic and seasonal patterns suggest that these associations reflect ecologically mediated cross-species transmission facilitated by mixed roosting, synchronized colony phenology, and spatial overlap. Our findings reveal greater permeability of coronavirus host boundaries than previously assumed and highlight multi-species bat colonies and ecological networks as key contexts for understanding and anticipating viral emergence.

## Introduction

Although the first coronavirus (CoV) was isolated in chicken embryos in 1937 (Beaudette et al. 1937), CoVs were not recognized as major human pathogens until the emergence of SARS-CoV in 2002-2003. Since the beginning of the 21^st^ century, five new coronaviruses capable of causing respiratory diseases in humans have been reported, including SARS-CoV, HCoV-NL63, HCoV-HKU1, MERS-CoV, and SARS-CoV-2 (Chen et al. 2020, Kesheh et al. 2022). These viral outbreaks have stimulated research into wildlife viruses and the remarkable diversity of coronaviruses in nature (Drexler et al. 2014, Anthony et al. 2017, Ar Gouilh et al. 2018, Latinne et al. 2024, Han et al. 2023). Recently, HKU5-CoV lineage 2was reported in *Pipistrellus* species. This virus efficiently uses human angiotensin-converting enzyme 2 (ACE2) as a functional receptor in bats and exhibits broad host tropism (Chen et al. 2025). Soriano-Tordera et al. (2025) found that a CoV obtained from Spanish bats can also use human ACE2. Although no human infection with these viruses have been reported to date, these findings highlight the relevance of bat coronaviruses to emerging diseases risk. Coronaviruses are large, enveloped viruses with single-stranded, positive-sense, non-segmented RNA genomes of approximately 30 Kb (Kipe & Howley 2013). They belong to the Coronaviridae family and are classified into four genera: *Alpha-*, *Beta-*, *Gamma-* and *Deltacoronavirus*, and two other putative genera *Epsilon-* and *Zetacoronavirus* (Zamperin et al. 2023, Hou et al. 2025). The seven CoVs reported in humans belong to αCoV and βCoV genera cause respiratory diseases that ranging from mild to severe. βCoVs include viruses that cause severe human diseases, such as *SARS-CoV*, *SARS-CoV-2* and *MERS-CoV*. Coronaviruses primarily exhibit respiratory and enteric tropism, although they may also display hepatic and neurological tropism in animals and humans. Recombination plays a major role in coronaviruses evolution and can facilitate the infection of new hosts and alter virulence (Graham & Baric 2010).

Bat evolution represents a highly successful and unique history among mammals, resulting in remarkable species diversity adapted to a wide range of environments. Bats are the second largest order of mammals, representing 22% of all extant mammalian species (https://www.mammaliandiversity.org). They have been implicated in numerous emerging infectious disease events and have been recognized as important reservoir hosts for viruses capable of crossing species barrier to infect humans and other domestic and wild mammals. Their role in viral diseases is well established, particularly for lyssaviruses and coronaviruses (Hayman et al. 2016, Han et al. 2023, Lantinne et al. 2024). Bats harbour the greatest known diversity of CoVs among mammals, and the α- and βCoV have been widely detected in bats across various regions worldwide (Drexler et al. 2014, Anthony et al. 2017, Wong et al. 2019, Zhou et al. 2022). Bats are largely asymptomatic to CoV infections (Aicher et al. 2022). Millions of years of bat evolution may have driven co-evolutionary processes between bats and viruses (Afelt et al. 2018). Bats possess a unique interferon system, which may help explain their ability to coexist with CoVs (Zhou et al. 2016, Morales et al. 2025). Bats have several unique features that may maximize their effectiveness as reservoir hosts of viruses. Their gregarious roosting behaviour, particularly during the summer season, where they closely cluster together, can significantly increase the likelihood of viral transmission within the same species and between different bat species (Serra-Cobo & López-Roig 2016). The higher frequency of interspecies transmission observed in the evolutionary history of coronaviruses, suggests that CoVs exhibit genus specificity rather than species specificity, even though they also exhibit a stronger association in those bat species that live sympatrically (Leopardi et al. 2018). Although cohabiting bat species do not always share the same CoVs, indicating the existence of some barrier to transmission between species (Anthony et al., 2013; Corman et al., 2013; Cui et al., 2007; Drexler et al., 2010; Smith et al., 2016; Tang et al., 2006), they also experience a greater probability of cross-species transmission.

In this study, we analysed bat CoV sequences obtained from field samples collected in different locations in Catalonia and the Balearic Islands, Spain. We performed a bipartite network analysis between bat species and the − and CoV subgenera, and we analysed the relationships between host and these viral groups. We also investigated how CoVs circulate and are transmitted among bat species, with particular emphasis on sarbecoviruses circulation within a multi-species bat colony. Finally, we analysed which bat species share CoV subgenera at the network and phylogenetic tree level and related them to the bat host ecology.

## Material and methods

### Ethics Statement

The animal study followed the ethical recommendations of the Spanish Legislation (Law 42/2007). Bat sampling was authorized by Department of Climate Action, Food and Rural Agenda of the Generalitat de Catalunya, the Conselh Generau d’Aran and the Conselleria de Medi Ambient i Territori of the Govern de les Illes Balears. All manipulations were performed avoiding pain and minimizing animal discomfort. The study was conducted in accordance with the ARRIVE guidelines (https://arriveguidelines.org).

### Sample collection

This study was conducted across 31 roosts located in Catalonia and the Balearic Islands (Spain) (Figure 1). Bats were randomly captured between 2020 and 2024. Sampling of bat colonies occurred year-round, excluding periods of hibernation and birthing.

**Figure 1.**
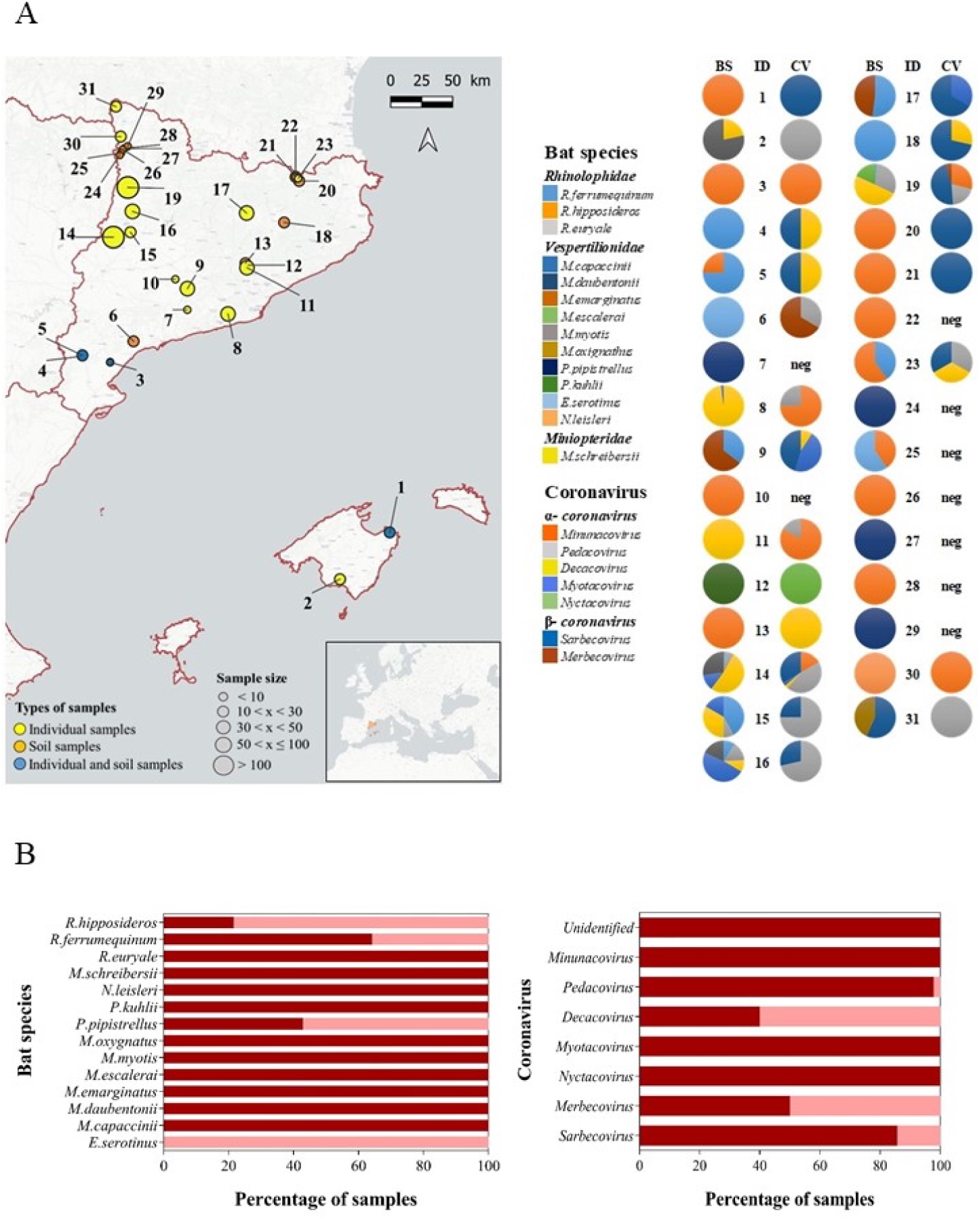
Overview of bat sampling and coronaviruses diversity across Catalonia and the Balearic Islands. **A:** Map of the sampled localities in Catalonia and the Balearic Islands. Pie charts indicate the proportion of bat species sampled (BS) and CoVs subgenera detected (CV) across the 31 localities (ID). Colours in the pie chart represent different species and CoVs, consistent with the colours of legend. **B:** Percentage of guano samples collected as individual faecal samples (red) and ground samples (pink) shown for bats and detected coronaviruses.

To interpret the interspecific circulation of viruses within a bat colony, it is essential to understand the colony’s dynamics. Accordingly, we analysed colony phenology alongside the interspecific circulation of coronaviruses (CoVs) in a multi-species bat colony, with specific focus on the circulation of sarbecoviruses.

Bats were sexed, identified to species, and placed inside a clean cotton bag during a short period to collect individual faeces. In some cases, faecal samples were obtained by placing clean sheets of aluminium foil under bat colonies. Each faecal sample was collected individually, except in a few cases where guano was pooled. All guano samples were obtained from monospecific colonies. All faecal samples were stored in sterilized vials containing RNAlater (Qiagen, Hilden, Germany) to preserve RNA integrity during the fieldwork and transport to the laboratory. Samples were frozen at - 80 °C before analysis.

### Sample preparation and detection of CoV RNA

The faecal suspension was clarified at 5,000 g for 10 min at 4 °C and total RNA was extracted from 150 μL supernatant using the Nucleospin Dx virus kit (Macherey-Nagel, Düren, Germany) following the manufacturer’s instructions. The reverse transcription (RT) step was performed using the SuperScript VILO cDNA Synthesis kit (Invitrogen, Carslbad, CA, USA) following the manufacturer’s instructions, using 14 μL of extracted RNA in a 20 μL total reaction volume. The first and the second PCR reactions (hemi-nested) were carried out using the Taq PCR Core kit (Qiagen, Hilden, Germany) following the manufacturer’s instructions, using pan-coronavirus primer set (PanCoV pol 15197/PanCoV pol 15635/PanCoV pol nested 15419) previously described (Ar Meriadeg et al. 2011, 2018). Five microliters of RT product were used in the first step PCR, in a 50 μL final reaction volume containing 50 pmol of the primer set PanCoV pol 15197/PanCoV pol 15635. For the second PCR reaction (hemi-nested), 5 μL of amplified product from the first PCR were added to a 50 μL reaction mixture containing 50 pmol of the second primer set (PanCoV pol 15635/PanCoV pol nested 15419). This methodology amplifies a conserved region of ORF1. After amplifications, 10 μL of the first PCR and the second (hemi-nested) PCR products were resolved on a 2% GelRed (Biotium)-stained agarose gel in TBE buffer and visualised under ultraviolet light. Amplicon sizes (440 and 220 bp for first and second PCR, respectively) were determined by comparing them with a 100-bp DNA ladder (Qiagen). Some *M. schreibersii* positive samples for CoVs were analysed two times to confirm the result. PCR products were purified using the PureLink PCR Purification kit (Invitrogen, Carlsbad, CA, USA) and sequenced bidirectionally using Sanger sequencing (Macrogen Inc., Madrid, Spain).

PCR amplicons of the positive samples of CoV from *M. schreibersii* obtained in Tremp were sequenced using Oxford Nanopore Technologies platform to gather information about sarbecoviruses circulation. The Nanopore Classic PCR tiling of SARS-CoV-2 virus protocol was followed with minor adjustments for adapting it to SQK-LSK109 kit. The obtained library was loaded in Flongle R9.41 MinION flow cells.

The bioinformatic analysis was conducted using the approach of Carcereny et al. (2024) and adapting it for the coronavirus analysis changing both the reference database and the read parameters (minimum of 100 bp and maximum of 700 bp). In brief, raw reads were trimmed, filtered, and clustered at 95% id. The resulting consensus sequence of each cluster was searched against custom reference database and those reads having a minimum BLAST id of 80% with any sequence in the database were kept. Finally, the coronavirus strains, the number of each hit and the depth-weighted BLAST id were recorded for each sample.

A negative control was sequenced and analysed along with the samples to assess for contamination and barcode bleeding. Neither was observed.

### DNA analysis of bat species

Given the relevance of the results obtained in the Tremp colony, the guano from the *M. schreibersii* individuals that tested positive for βCoV was analysed to confirm the bat species. The guano preserved in RNAlater was vortexed for 1 min and clarified at 5,000 g for min at 4 °C. Total DNA was extracted using a QIAmp Fast DNA Stool Mini Kit (Qiagen) following the manufacturer’s instructions. A 1,643 bp fragment of the entire mitochondrial *Cytochrome b* and partial *D-loop* gene was amplified using the primer set (Chiro_cytb521+: 5’-TGGCATGAAAAATCACCGTTGT-3’ / Chiro_cytb1743-: 5’-ATGGCCCTGAAGAAAGAACCAGATG-3’) as previously described (Puechmaille et al. 2011).

The extracted DNA was subjected to PCR using a Taq PCR Core kit (Qiagen) following manufacturer’s instructions. Five microliters of DNA were added to 10 pmol of the primer set (Chiro_cytb521+ / Chiro_cytb1743-) in a final volume of 25 μL with the following amplification conditions: an initial denaturation step at 94 °C for 2 min, then 10 cycles at 94 °C for 30 s, 65 °C for 30 s (with a reduction of 2 °C every 2 cycles), 72 °C for 40 s, followed by 30 PCR cycles at 94 °C for 20s, 55 °C for 30 s, 72 °C for 40 s and a final elongation step at 72 °C for 3 min.

After amplification, 10 μL of the PCR products were visualised on a 2% GelRed (Biotium Hayward)-stained agarose gel in TBE buffer and then compared with a 1 Kb DNA ladder (NEB). Positive samples with the appropriate amplicon size were purified using the PureLink PCR kit (Invitrogen) and bidirectionally sequenced (Macrogen) using the PCR primers mentioned earlier.

### Phylogenetic analysis

Two phylogenetic trees were constructed using the positive sequences obtained in Catalonia and the Balearic Islands: one based on αCoV sequences and the other on βCoV sequences. For the αCoV and βCoV trees we designed a pipeline capable of dealing with hundreds of sequences. First all the reverse strand sequences were reverse-complemented using revcomp (v1.9c). Then all the sequences belonging to the same bat_ID were aligned using MAFFT (v7.475) G-INS-I (Katoh et al. 2002) and feed to HMMER (v3.3.2) to generate a consensus with hmmbuild and hmmemit. The consensus sequence for each batID strand was examined using bl2seq (v2.11.0) (Altschul et al. 1990) and homogenized if necessary, using revcomp (v1.9c). After a rough classification into alphacoronavirus or betacoronavirus using blastn (v2.11.0) (Altschul et al. 1990), an alignment for each group was performed using MAFFT (v7.475) L-INS-I (Katoh et al. 2002) adding two sequences of the other group to act as a root. The resulting alignments were examined using Jalview (v2.11.4.1) (Waterhouse et al. 2009) and trimmed to the core conserved region in each case. The trimmed alignment was processed by IQ-Tree webserver (Trifinopoulos et al. 2016) with the default parameters except for “number of bootstrap alignments” set to 2500; “Maximum Iterations” set to 2000; “SH-aLRT branchtest #replicates” set to 2500; “Approximate Bayes test” set to “yes”.

A more personalized approach was used for generating the Tremp cave tree due to the novelty of the results obtained. Quality trimming, assembling and consensus generations were performed using Bioedit sequence alignment editor software (v7.2.5) (Hall 1999). Alignment of original consensus sequences, with CoV sequences obtained in Genbank were performed using MAFFT (v7.310) L-INS-I (Katoh et al. 2002). The best substitution Model was identified by FindModel (Posada and Crandall 1998). Phylogenetic trees were conducted with MEGA11 software (Tamura et al. 2021) using Maximum likelihood models with 1,000 bootstraps. The resulting trees were visualized using FigTree (v1.4.4) and edited for readability using Illustrator (v28.1.1).

Species diversity and spatio-temporal community composition of Tremp locality was performed by alluvial graph from library Alluvial in R.

### Network analysis

We built a dataset of bat species-coronavirus associations from 31 roost sampled during July 2020 to March 2024 in Catalonia and Balearic Islands (Figure 1). We used this data to construct a weighted bipartite undirected network between 13 bat species and 7 subgenera coronaviruses. Bat species with no positive samples were excluded from the dataset. Network links were weighted and defined as percentage of positives samples relative to the total number of samples for each bat species and subgenus of coronavirus.

From this network, we calculated four structural metrics that describe different aspects of network assembly and distribution of interactions. First, weighted modularity (Qw), calculated by the DIRTLPAwb + algorithm. A modular network comprises subgroups of species in which interactions are stronger and more frequent than species out of these subgroups, which may reveal functional groups in the network. Qw varies from zero to one, the latter representing a perfectly modular network. Second, complementary specialization through the H2′ metric. It quantifies how unique, on average, are the interactions made by species in the network, considering interaction weights and correcting for network size. It varies from zero to one, the latter corresponding to a specialized network where interactions perfectly complement each other because species do not share partners. The index was normalized to vary from zero to one, with one representing a perfectly nested network. Metric significance (Qw, H2′, Connectance and WNODA) was assessed using Null model simulations conducted by the *r2dtable* algorithm to preserve marginal totals. For each index, 1,000 randomized matrices were generated with *nullmodel()*, and the observed value was compared to the null distribution to obtain one-tailed p-values. The significance threshold was considered *p* ≤ 0.05.

To assess the topological importance of individual nodes, we computed several centrality measures of undirected weighted bipartite network, where edge weights represented the percentage of positives. We measured degree (interaction strength), betweenness and closeness centralities, eigenvector centrality (to identify nodes connected to other well-connected nodes) and edge-level betweenness (using the inverse of edge weights). These values were then aggregated per node to derive total edge betweenness (sum of all associated edges), mean edge betweenness, and the number of incident edges.

We also used the single-mode projection of bat nodes as it provides a more generalized picture of how bat species can be associated according to the coronaviruses they share. In this case the network links were also weighted. A weighted Jaccard similarity matrix was also computed to quantify the proportion of shared viruses relative to the total combined viral associations between each pair of species. On this single-mode projection, weighted centrality metrics were recalculated. Weighted degree was defined as the sum of edge weights per node. Betweenness and closeness centralities were computed using the inverse of similarity weights, treating stronger similarities as shorter topological distances. Eigenvector centrality was also obtained to evaluate a node’s influence through its connected neighbours. The virus frequency per species from the original bipartite network was included, and a weighted centrality index was defined as the product of weighted degree and virus frequency. All centrality metrics were normalized between 0 and 1, and a composite relevance index was calculated as the mean of the normalized values to rank species according to their structural role in the network. To identify global patterns and relationships among the centrality measures, a Principal Component Analysis (PCA) was performed. All centrality metrics were linearly normalized (min–max scaling) between 0 and 1.

Finally, roost-share bat associations weighted network was performed to determine which bat species were solitary or roosting with other bat species. We used Louvain community analysis to identify species associations according to whether they share refuge and dendrogram view community was also performed.

All networks were exported to Cytoscape (v3.10.3) in GraphML format for visualization (Shannon et al. 2003).

## Results

### CoV-RNA analysis

We obtained 835 samples of which 712 were individual samples from bat captures and 123 were ground samples. The analysed samples were obtained from 31 different roosts with a total of 14 bat species corresponding to 3 families. Of the 31 locations surveyed, only 9 (29%) tested negative for CoV and only one of these corresponded to a multi-species colony (Figure 1).

Of the 835 samples analysed, 18.92% tested positive for CoV RNA, including 11.50% positive for αCoV and 7.42% for βCoV, relative to the total samples (Supplementary table S1).

All positive samples were identified at the subgenera level except 2 of them which were identified at the genera level.

We identified 5 subgenera of αCoV and 2 subgenera of βCoV (Supplementary table S1, Figures 1 and 2). Figure 3 shows the phylogenetic trees for αCoV and βCoV constructed with the positive sequences obtained in this work. The most frequently detected subgenus among αCoVs has been *Pedacovirus* (47.87% of αCoVs subgenera identified), while *Sarbecovirus* (SARS-like-CoV) has been the dominant βCoV (93.55%). We found the greatest diversity of CoVs in *R. euryale*, *R. hipposideros*, *R. ferrumequinum* and *M. schreibersii*.

**Figure 2.**
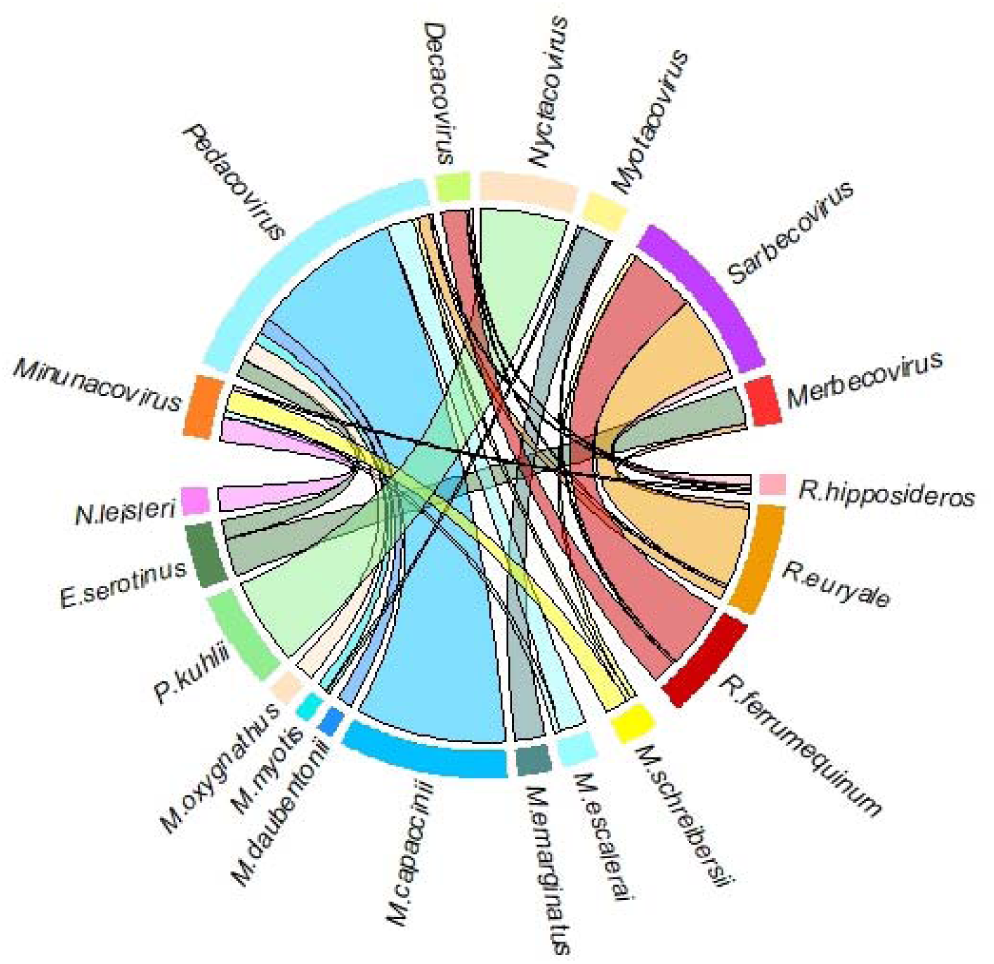
Chord Diagram network between coronavirus subgenera and bat species sampled. Sector size is proportional to the total number of positive detections for each bat species or coronavirus subgenus. Chords represent host–virus associations, and their width is proportional to the percentage of positive samples for each interaction. Colours identify bat species and coronavirus subgenera and are consistent with those used throughout the manuscript. The network summarizes the distribution of coronavirus subgenera across bat hosts and illustrates shared viral associations among species.

**Figure 3.**
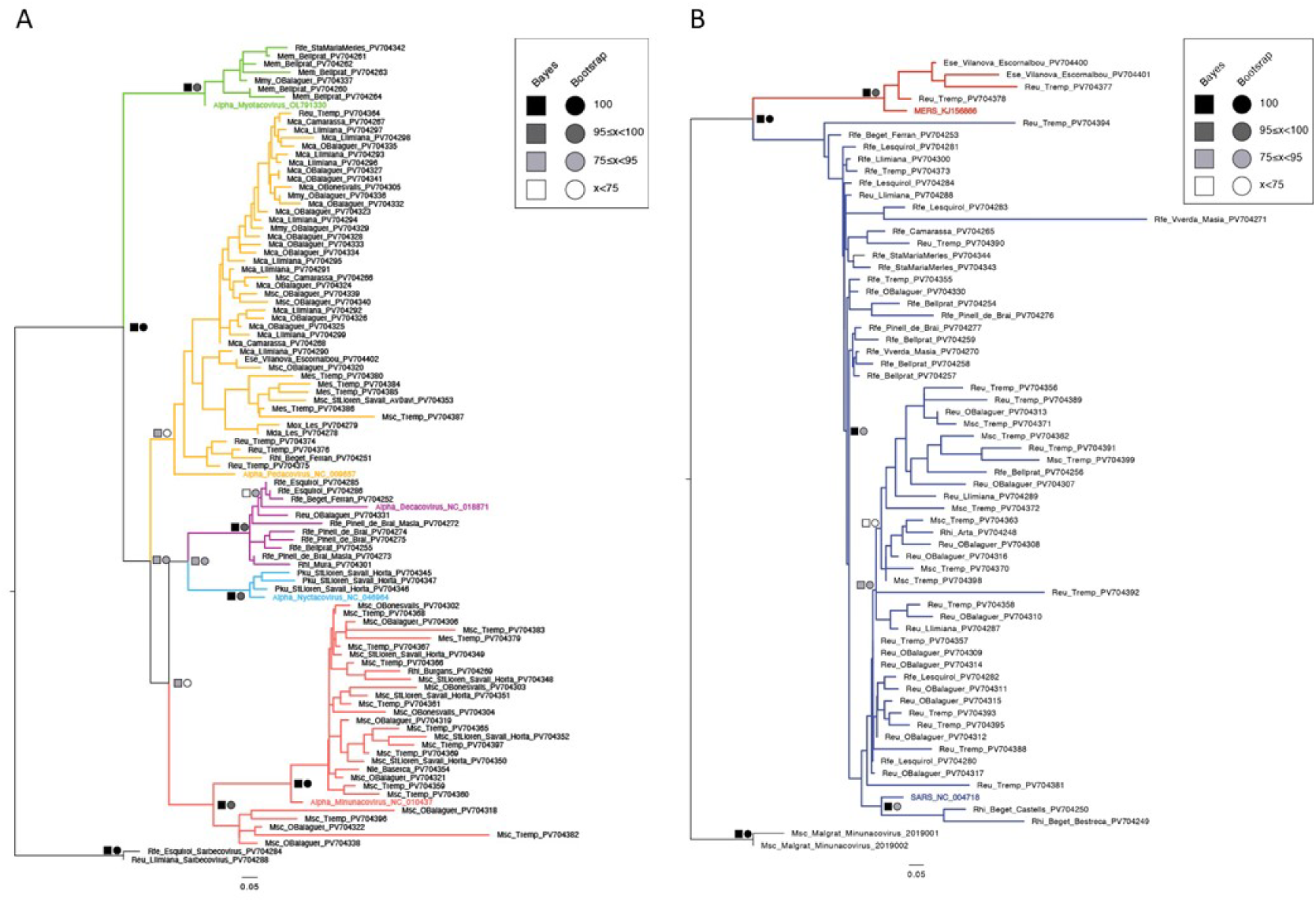
Maximum-likelihood phylogenetic trees based on partial RdRp gene sequences. The trees show (A) αCoV and (B) βCoV identified in bat guano samples collected in Catalonia and the Balearic Islands. The phylogenies reveal the diversity of circulating coronavirus lineages and their relationships with previously described bat coronaviruses, including the clustering of the newly detected *Sarbecovirus* and *Merbecovirus* sequences within established βCoV lineages.

### Interspecific circulation of CoV in a multi-species colony

Over the course of four years, 221 bats of four species and 3 families were sampled in Tremp locality. All four bat species were clearly identified and corresponded to *M. schreibersii*, *R. euryale*, *R. ferrumequinum* and *M. escalerai*. The first two species are the most abundant and have a similar phenology. In spring, both species form colonies that can exceed 600 individuals (May) at their peak, although their number varies annually. In autumn, both species are also present but in smaller numbers. The other two species are less abundant and less frequent (Table S2, Figure 4).

**Figure 4.**
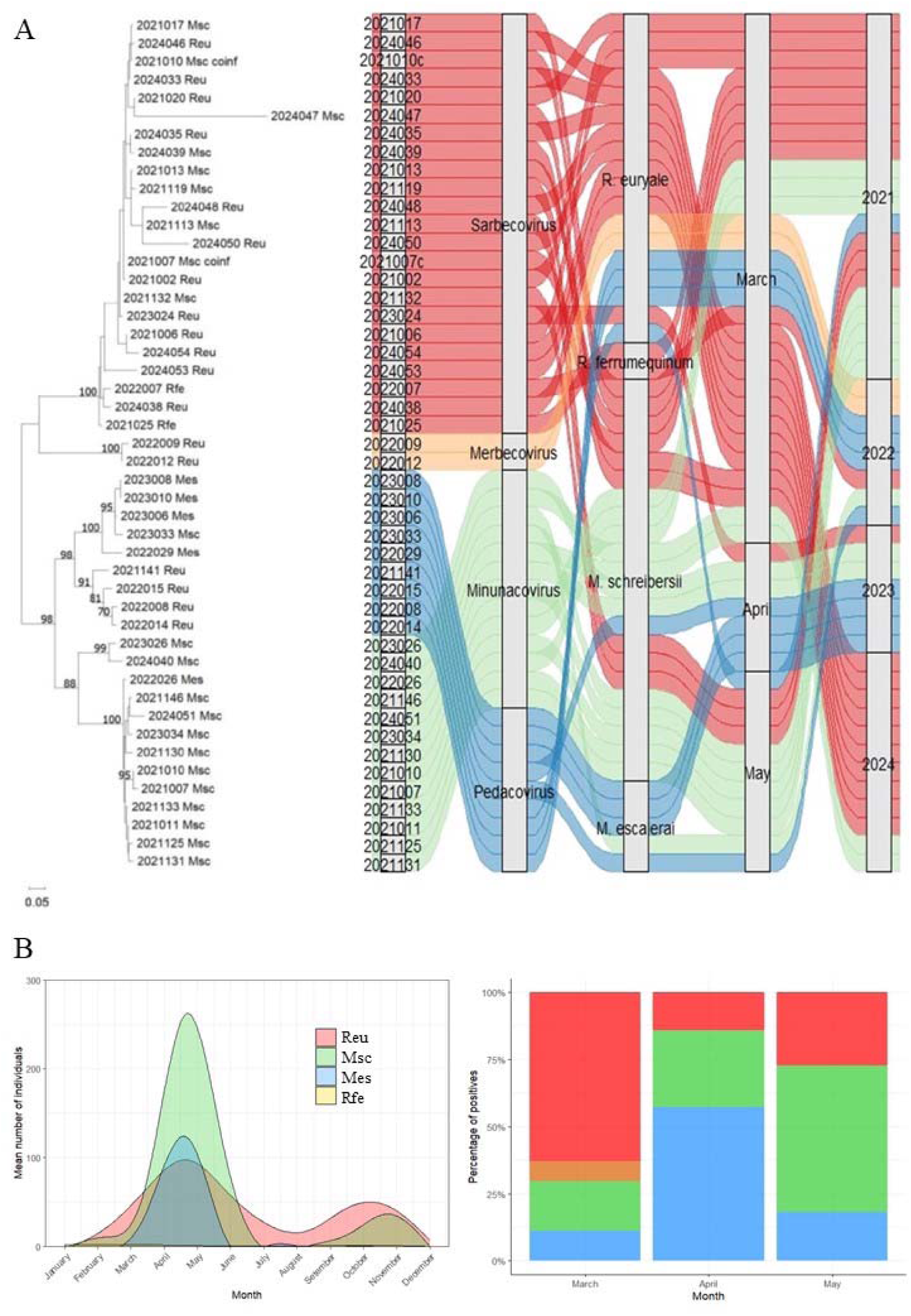
Seasonal host turnover shapes coronaviruses circulation within a multi-species bat colony. **A:** Phylogenetic tree (left side) and alluvial diagram showing the temporal dynamics and relationship between CoVs and the bat species sampled (right side). The acronym coinf at the end of the sequences indicates coinfection of the αCoV and βCoV in the same bat. **B:** Temporal dynamics of the multi-species bat colony (left side) and CoVs (right side) in the Tremp cave. The abbreviated names and colors of the coronaviruses correspond to those in subfigure A.

The diversity and composition of the coronaviruses found varied according to seasonality. βCoVs were predominant during the month of March, especially *Sarbecovirus* (63%), while αCoVs (*Minunacovirus* and *Pedacovirus*) did not exceed 30%. Subsequently, with the arrival of a greater number of *M. schreibersii*, the αCoV detected increased to 86%.

CoV RNA detection and characterization enabled the identification of the sarbecoviruses to be found for the first time in *M. schreibersii*. Seven bats of this species have tested positive for SARS-like-CoV. On the other hand, the analysis done with the MinION confirmed the coinfection of SARS-like-CoV and αCoVs (*Minunacovirus*) in two more *M. schreibersii*. DNA extracted from guano samples that tested positive for a βCoV in *M. schreibersii* was subsequently analysed to confirm the bat species. Additionally, this study reports the first identification of a *Merbecovirus* in *R. euryale* in the same colony (Figure 4, Supplementary table S2).

We identified two αCoV subgenera (*Minunacovirus* and *Pedacovirus*) and two βCoV subgenera (*Sarbecovirus* and *Merbecovirus*) circulating in this shelter (Figure 4). As the alluvial diagram reveals, βCoVs circulate in *M. schreibersii* throughout all months and years sampled.

### Bat–coronavirus network structure

Regarding general network descriptors, our weighted bipartite network comprises 20 nodes and 26 edges, with an average of 2.6 neighbours per node (Figure 5A). The network density (0.137) and heterogeneity (0.732) indicated a heterogeneous distribution of interactions, while the centrality value (0.337) suggests that a subset of species occupies relatively more central positions within the network. The network structure was also assessed using the four calculated structural metrics: complementary specialization (H2′ = 0.795, p < 0.001), weighted modularity (Qw = 0.648, p < 0.001), nestedness (wNODF = 8.67, p = 0.041) and connectivity (0.286, p < 0.001), indicating a highly structured network and a non-random interaction pattern.

**Figure 5.**
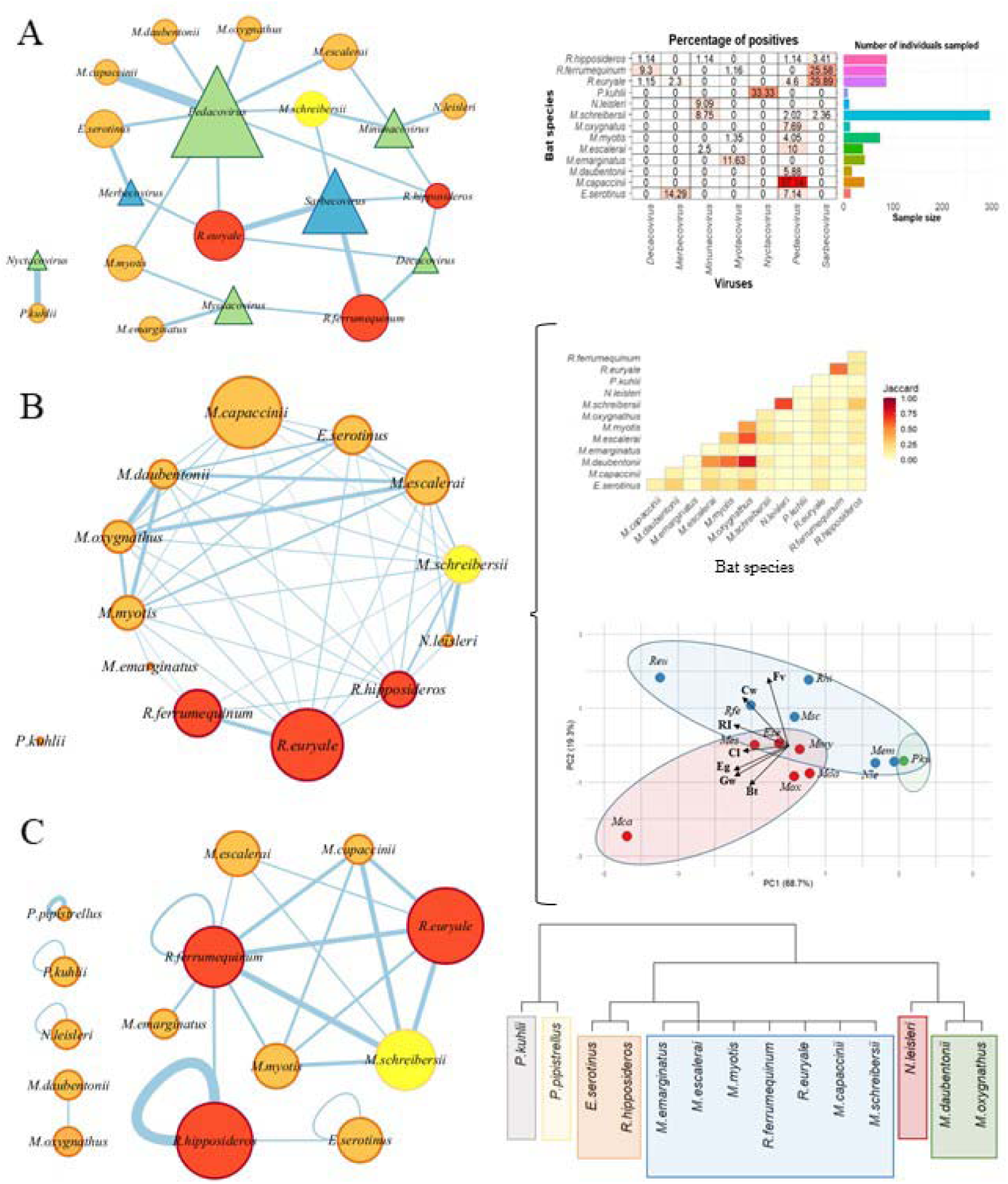
Network analyses identify *Rhinolophus* and *Miniopterus* bats as central connectors of coronavirus circulation. **A:** weighted bipartite network and percentage of positive samples for each bat species and CoVs subgenus. **B:** single projection matrix and Jaccard matrix between bats species (upper right) and PCA (lower right). **C:** Roost- sharing bat species network and cluster tree according to their compartmentalization. Bat species: *Ese*, *E. serotinus*; *Mca*, *M. capaccinii*; *Mda*, *M. daubentonii*; *Mem*, *M. emarginatus*; *Mes*, *M. escalerai*; *Mmy*, *M. myotis*; *Mox*, *M. oxygnathus*; *Msc*, *M. schreibersii*; *Nle*, *N. leisleri*; *Pku*, *P. kuhlii*; *Ppi*, *P. pipistrellus*; *Reu*, *R. euryale*; *Rfe*, *R. ferrumequinum*; *Rhi*, *R. hipposideros*.

Analysis of node centrality metrics revealed notable heterogeneity in the role of bat species and viral taxa within the network (Table 1, Figure 5A). Among bat hosts, *R. euryale* and *R. ferrumequinum* displayed the highest betweenness centrality values (32 and 28, respectively), suggesting they act as important connectors between otherwise separate parts of the network. In contrast, *Myotis capaccinii* and *Eptesicus serotinus* showed high degree centrality (57.14 and 21.43, respectively), indicating numerous interactions, although their roles as intermediaries were limited. *P. kuhlii* and *Nyctacovirus* formed an isolated dyad with unusually high closeness centrality values (33.33), reflecting their exclusive connection in a separate component of the network (Table 1). In terms of eigenvector centrality, *M. capaccinii* (0.957) stood out as the most influential bat species in terms of connection to other well-connected nodes, followed by *M. escalerai* (0.168) and *E. serotinus* (0.128). This suggests that *M. capaccinii*, despite limited betweenness, is topologically embedded in a highly interactive network region.

**Table 1.** Weighted centrality metrics for bat species and coronavirus subgenera in the bipartite host–virus interaction network. Centrality measures calculated from the weighted bipartite network describing associations between bat hosts and coronavirus (CoV) subgenera detected in guano samples collected from bat roosts in Catalonia and the Balearic Islands, Spain, during 2020–2024. Edge weights represent the percentage of positive samples for each bat species–virus subgenus association relative to the total number of samples for that bat species. Bat species: *Ese*, *E. serotinus*; *Mca*, *M. capaccinii*; *Mda*, *M. daubentonii*; *Mem*, *M. emarginatus*; *Mes*, *M. escalerai*; *Mmy*, *M. myotis*; *Mox*, *M. oxygnathus*; *Msc*, *M. schreibersii*; *Nle*, *N. leisleri*; *Pku*, *P. kuhlii*; *Reu*, *R. euryale*; *Rfe*, *R. ferrumequinum*; *Rhi*, *R. hipposideros*.

| Bat species | Centrality |  |  |  | Edge_betweenness centrality |  |  |
| --- | --- | --- | --- | --- | --- | --- | --- |
|  | Degree | Betweenness | Closeness | Eigenvector | Total_edge_betweenness | Mean_edge_betweenness | N_edges |
| *Ese* | 21.43 | 16 | 0.120 | 0.128 | 49 | 24.50 | 2 |
| *Mca* | 57.14 | 0 | 0.150 | 0.957 | 17 | 17.00 | 1 |
| *Mda* | 5.88 | 0 | 0.110 | 0.098 | 17 | 17.00 | 1 |
| *Mem* | 11.63 | 0 | 0.052 | 0.000 | 17 | 17.00 | 1 |
| *Mes* | 12.50 | 14 | 0.133 | 0.168 | 45 | 22.50 | 2 |
| *Mmy* | 5.40 | 14 | 0.107 | 0.068 | 45 | 22.50 | 2 |
| *Mox* | 7.69 | 0 | 0.118 | 0.129 | 17 | 17.00 | 1 |
| *Msc* | 13.13 | 14 | 0.094 | 0.039 | 45 | 15.00 | 3 |
| *Nle* | 9.09 | 0 | 0.077 | 0.002 | 17 | 17.00 | 1 |
| *Pku* | 33.33 | 0 | 33.330 | 0.000 | 1 | 1.00 | 1 |
| *Reu* | 37.94 | 32 | 0.142 | 0.116 | 81 | 20.20 | 4 |
| *Rfe* | 36.04 | 28 | 0.135 | 0.034 | 73 | 24.30 | 3 |
| *Rhi* | 6.83 | 0 | 0.085 | 0.024 | 17 | 4.25 | 4 |
| **Subgenus coronavirus** |  |  |  |  |  |  |  |
| *Merbecovirus* | 16.59 | 0 | 0.106 | 0.035 | 17 | 8.5 | 2 |
| *Pedacovirus* | 99.66 | 85 | 0.157 | 1.000 | 187 | 20.8 | 9 |
| *Myotacovirus* | 14.14 | 16 | 0.057 | 0.002 | 49 | 16.3 | 3 |
| *Minunacovirus* | 21.48 | 17 | 0.089 | 0.013 | 51 | 12.8 | 4 |
| *Decacovirus* | 11.59 | 0 | 0.109 | 0.008 | 17 | 5.67 | 3 |
| *Sarbecovirus* | 61.24 | 51 | 0.142 | 0.075 | 119 | 29.8 | 4 |
| *Nyctacovirus* | 33.33 | 0 | 33.330 | 0.000 | 1 | 1.0 | 1 |

Viral taxa showed a similar degree of structural asymmetry: *Pedacovirus* exhibited the highest centrality values across all metrics, including degree (99.66), betweenness (85), and eigenvector centrality (1.00), confirming its dominant and integrative role in the network. *Sarbecovirus* and *Minunacovirus* also showed high betweenness (51 and 17, respectively), suggesting their involvement in cross-host transmission pathways. By contrast, *Decacovirus* and *Nyctacovirus* exhibited low centrality scores, consistent with a more peripheral or specialized ecological position. Overall, these centrality measures highlight the existence of key host species and viral subgenera that disproportionately shape the structure and potential connectivity of the bat–coronavirus interaction network.

The weighted centrality analysis allowed for the identification of bat species potentially more influential in the viral sharing network. Species from the *Rhinolophus* genus ranked among the most prominent: *R. euryale* showed the highest centrality, with a high viral frequency (4 viruses) and elevated closeness and eigenvector values, indicating a central position in the network. Similarly, *R. ferrumequinum* achieved a high value, derived from a moderate degree but a high viral frequency (3 viruses). Surprisingly, *M. capaccinii* ranked as the third most relevant species, despite being associated with only one virus (*Pedacovirus*); this result reflects an exceptionally high weighted degree, along with maximum scores in closeness, betweenness, and eigenvector, suggesting it occupies a strategic position within the network. Species such as *M. escalerai*, *E. serotinus*, *R. hipposideros*, and *M. schreibersii* also exhibited high weighted centrality values, particularly the first two for their connectivity and the latter two for their higher viral frequency. In contrast, species like *M. emarginatus*, *N. leisleri*, and *P. kuhlii* showed the lowest values across all metrics, with *P. kuhlii* being entirely isolated within the network (weighted centrality = 0). These results point to the existence of a core group of species centrally located in the viral interaction network, which could play a key epidemiological role, either as potential transmission bridges or as reservoirs with strong ecological or phylogenetic connections.

### Single-mode projection network

The weighted similarity analysis using the Jaccard index revealed clear patterns in virus sharing among bat species. High similarity values were detected between species of the *Myotis* genus, notably the pair *M. daubentonii*–*M. oxygnathus* (index = 0.765), followed by *M. daubentonii*–*M. myotis* (0.560) and *M. escalerai*–*M. oxygnathus* (0.615), suggesting a high overlap in the set of shared viruses. In contrast, species such as *P. kuhlii* and *N. leisleri* showed practically zero similarity with most other species, indicating a probable viral specialization or isolation. Notable relationships were also identified between species of different genera, such as *E. serotinus* and *M. oxygnathus* (0.325). A high shared exposure to common virus sources was also observed among the Rhinolophidae family, particularly between *R. euryale* and *R. ferrumequinum* (0.566) (Figure 5B).

Community analysis on the unimodal projection (species × species network) revealed the existence of three main groups of bats, structured according to virus-sharing patterns and connection intensity (Figure 5B). The first group mainly includes species from the *Myotis* genus, such as *M. capaccinii*, *M. escalerai*, *M. daubentonii*, and *M. oxygnathus*, characterized by high Jaccard similarity values and centrality, suggesting high ecological and/or phylogenetic cohesion. The second group is primarily composed of species from the *Rhinolophus* genus (*R. euryale*, *R. ferrumequinum*, *R. hipposideros*) and associated species, such as *M. schreibersii* and *Myotis emarginatus*, which share several viruses among themselves but show fewer connections to other groups. Finally, the third, much more peripheral, group includes species like *P. kuhlii*, with marginal participation in the network and no significant connections (Figure 5B).

### Roost-sharing bat species network

The analysis of the network of species sharing refuges showed a greater relationship between the two species of *Rhinolophus* (*R. ferrumequinum* and *R. euryale*) with mainly *M. schreibersii*, *M. capaccinii* and *M. myotis* and to a lesser extent with *M. emarginatus* and *M. escalerai*. Virtually all other species were solitary or shared refuges occasionally (Figure 5C). These results indicate that the community structure of the network reflects not only the number of shared viruses but also co-occurrence patterns and weighted connectivity, likely influenced by ecological and habitat-use factors.

## Discussion

### Structured viral communities and ecological constraints on CoVs circulation

Our results reveal that CoVs circulation in European bats is not random but instead highly structured, as demonstrated by the significant modularity and specialization observed in the virus–host interaction network. These findings align with previous studies suggesting that ecological and evolutionary processes jointly shape host–virus associations in bats (Drexler et al. 2014; Leopardi et al. 2018). The bat species tend to form distinct ecological clusters in which viral exchange appears to be more frequent.

Such structure likely reflects underlying ecological constraints, including roosting behaviour, phylogenetic relatedness, and habitat overlap.

A detailed analysis of αCoV showed that the subgenera *Decacovirus* and *Nyctacovirus* were primarily associated with a single bat genus—*Rhinolophus* and *Pipistrellus*, respectively—whereas *Minunacovirus*, *Myotacovirus*, and particularly *Pedacovirus* were detected in multiple genera. These findings are consistent with the αCoV phylogeny reconstructed by Latinne et al. (2024). Notably, *Rhinolophus* species harboured all αCoV subgenera identified in this study (except *Nyctacovirus*, which was restricted to *P. kuhlii)*. The *Minunacovirus* subgenus, originating from miniopterid bats, was detected in several bat species across three different families, suggesting that cross-species transmission events have occurred. βCoV analysis showed that *Rhinolophus* species were the main carriers, particularly of sarbecoviruses.

While specialization (H2′) suggests relatively restricted interaction patterns at the global network level, the existence of non-zero connectance and nestedness indicates that complete isolation between host groups does not occur. Instead, a subset of more generalist or central species contributes disproportionately to the maintenance of viral diversity across the network. This dual pattern (specialized modules interconnected by key species) provides an important ecological framework for understanding how coronaviruses persist and evolve within bat communities.

### Cross-species transmission in multi-species colonies: ecological drivers

A central contribution of this study is the identification of ecologically mediated cross-species transmission in multi-species bat colonies. Phylogenetic clustering of viral sequences, together with temporal overlap in infection patterns, is consistent with the interpretation that the novel host associations do not appear to be the result of long-term co-divergence alone, but rather of ongoing interspecific transmission processes.

Tremp multi-species colony provides compelling evidence that mixed roosting conditions facilitate viral exchange across host boundaries. The locality hosts *R. euryale* and *M. schreibersii*, which usually form a mixed colony from March to May. Phylogenetic analyses reveal that SARS-like-CoV sequences obtained from *M. schreibersii* cluster closely with those from *R. euryale*, suggesting possible transmission from *R. euryale* to *M. schreibersii*. We detected the SARS-like-CoV in *M. schreibersii* in March and May 2021 and in March 2024, but not in 2022 and 2023, a pattern consistent with the colony’s phenology. *R. euryale* usually arrives in early March, and *M. schreibersii* begins arriving in late March when the *R. euryale* colony is established. In 2022, we did not find *R. euryale* or *M. schreibersii* with sarbecoviruses, and by April 2023, unlike other years, *R. euryale* had not formed a colony in the refuge.

The synchronization of colony phenology, especially during maternity periods, likely enhances contact rates between species and promotes viral transmission. Shared maternity colonies, characterized by high densities and physiological stress, further intensify opportunities for cross-species infection by increasing direct contact rates or elevating the likelihood of indirect exposure via faeces, urine, or aerosols. These findings are consistent with prior observations that bat aggregation and high-density roosting increase opportunities for pathogen spillover (Serra-Cobo & López-Roig 2016). Moreover, the detection of shared viral lineages among phylogenetically distant hosts indicates that ecological overlap may outweigh phylogenetic constraints.

### Expansion of βCoVs host range: implications for viral evolution

To date, species of the family Rhinolophidae have repeatedly been identified as reservoirs of sarbecoviruses. Our results confirm for the first time with certainty the presence of sarbecoviruses in *M. schreibersii* (family Miniopteridae) and document the presence of *Merbecovirus* in *R. euryale*. Moraga-Fernández et al. (2022) reported preliminary evidence, but they could not confirm the presence of sarbecoviruses in *M. schreibersii*.

This represents a significant expansion of the known host range of these βCoVs and suggests that host-switching events may be more frequent than previously assumed. The presence of sarbecoviruses in *Miniopterus fuliginosus* (Zhang et al. 2025), recently confirmed in China, is consistent with what has just been reported. Similarly, the detection of *Merbecovirus* in *R. euryale* is unexpected given that this subgenus has historically been restricted to vespertilionid bats. The high similarity of the sequences suggests that MERS-like-CoVs may have a broader host range than previously recognized, expanding the potential interface between viral groups of known zoonotic relevance.

From an evolutionary perspective, the presence of closely related viral sequences across different host taxa supports the hypothesis that recombination and host-jumping play a central role in CoVs diversification (Graham & Baric 2010). The coexistence of multiple CoVs lineages within the same ecological network increases the likelihood of coinfections, providing opportunities for genetic exchange and the emergence of novel variants. Furthermore, *M. schreibersii* is a migratory species that can undertake seasonal movements exceeding 300 km (Hutterer et al. 2005; Serra-Cobo et al. 1998), and may contribute to the regional dissemination of viral diversity (Pons-Salort et al. 2014, Colombi et al. 2019) (Figure S1).

Importantly, recent evidence demonstrating that certain bat CoVs can utilize human ACE2 receptors (Chen et al. 2025; Soriano-Tordera et al. 2025) underscores the relevance of expanding host ranges in assessing zoonotic risk. Our findings suggest that ecological scenarios facilitating inter-species transmission could also create conditions conducive to the emergence of viruses with broader host tropism.

### Central connector species and their role in viral flow

Network centrality analyses identified bat species, particularly within the genera *Rhinolophus* and *Miniopterus*, as key connectors within the virus–host network. Among hosts, *R. euryale* and *R. ferrumequinum* show the highest centrality scores, suggesting that they act as important connectors between otherwise weakly connected modules. These results agree with previous studies conducted in China indicating that Rhinolophidae play a central role in the cross-species transmission of CoVs, contributing disproportionately to host switching events for αCoVs and serving as major donors for βCoVs (Latinne et al. 2024).

Interestingly, *M. schreibersii* also ranks among the most central taxa, particularly in the unimodal bat–bat network, where it forms strong connections with *Rhinolophus* and other sympatric genera. Although traditionally regarded as a host predominantly associated with αCoVs, its network position indicates that it acts as a bridging species capable of facilitating viral flow between otherwise segregated host clusters. The detection of a *Sarbecovirus* in *Miniopterus* is consistent with its inferred structural role and supports the hypothesis that central species may be disproportionately prone to acquiring novel viral lineages through ecological contact.

In our study, species with high degree, betweenness, and eigenvector centrality were also associated with higher viral diversity, suggesting that ecological traits such as large colony sizes, high roost fidelity, and extensive spatial overlap with other species enhance their capacity to accumulate and redistribute viral diversity. The identification of these connector species has important implications for surveillance strategies. Monitoring efforts that focus on highly central bat species may provide more efficient early detection of novel CoVs and transmission events than approaches based solely on species richness or abundance.

### Rethinking host specificity in bat CoVs

Traditionally, CoVs in bats have been considered to exhibit a relatively high degree of host specificity, often constrained at the genus or family level. While previous studies have acknowledged occasional host-switching events, these have generally been viewed as exceptions rather than a defining feature of CoV ecology.

Our findings challenge this prevailing paradigm. The combined evidence from network topology, phylogenetic analyses, and ecological observations indicates that host boundaries are more permeable than previously assumed. Rather than being strictly compartmentalized, bat CoV communities appear to function as dynamic systems in which ecological interactions facilitate periodic exchanges between host species.

This shift in perspective has important implications for understanding CoV evolution and emergence. It suggests that the potential for cross-species transmission is an intrinsic property of bat CoV systems, modulated by ecological context rather than strictly limited by host taxonomy. These results highlight the need to reconsider host range models of CoVs and underscore the epidemiological importance of monitoring multi-species bat colonies to better anticipate the evolutionary trajectories of CoVs and identify preventive measures.

## Conclusion

In summary, our study provides integrative evidence that CoV circulation in European bats is shaped by a combination of ecological structure and inter-species transmission processes. The detection of expanded host ranges for key βCoVs subgenera, together with the identification of central connector species and ecologically driven transmission pathways, reveals a more dynamic and interconnected viral landscape than previously recognized. Ultimately, our work challenges the prevailing paradigm of strict coronavirus host specificity in bats, demonstrating that host boundaries are more permeable than previously assumed, and highlighting the critical role of ecological interactions in shaping viral diversity and emergence.

## Data availability

The datasets generated and/or analysed during the current study are available from GenBank data base (PV704248-PV704402).

## Acknowledgements

We wish to thank the Departament de Medi Anmbient i Sostenibilitat of the Generalitat de Catalunya, the Departament de Medi Ambient of the Conselh Generau d’Aran), the National Park of Aigu estortes i Estany de Sant Maurici (Lleida, Spain), the Conselleria de Medi Ambient, Agricultura i Pesca of the Govern de les Illes Balears (Spain), the Natural Park of Sant Llorenç del Munt i l’Obac (Barcelona, Spain), the Parc of Garraf (Barcelona, Spain), the Malgrat de Mar City Council, Marina Berthet Garayoa and Marta Bordas for their contributions in facilitating the fieldwork logistics. We also thank Arnau Serra for his valuable contribution during the fieldwork.

## Ethics

The study followed the ethical recommendations of the Spanish Legislation (Law 42/2007) and was approved by the Bioethics Committee of the University of Barcelona. Bat sampling in Catalonia was authorized by the Departament d’Acció Climàtica, Alimentació i Agenda Rural of the Generalitat de Catalunya and under permit of the Natural Park of Sant Llorenc del Munt and Serra de l’Obac. In Menorca, the bat capture permits were granted by the Conselleria de Medi Ambient i Territori of the Govern de les Illes Balears. All procedures were conducted to minimize pain and animal discomfort. The study was conducted in accordance with the ARRIVE guidelines (https://arriveguidelines.org).

## Author Contributions

**Jordi Serra-Cobo**: Conceptualization, Fieldwork, Methodology, Supervision, Writing – review & editing. **Abir Monastiri**: Responsible for coronavirus analysis, DNA analysis, Writing & review. **Carlos Herrera-Ubeda**: Phylogenetic analysis, Writing & review. **Nuri Llorens Bàrbara:** Phylogenetic analysis, Writing & review. **Maria I Costafreda**: Coronavirus analysis, Writing & review. **Albert Carcereny**: Coronavirus analysis, Writing & review. **Marc López-Roig:** Conceptualization, Fieldwork, Network analysis, Writing – review & editing. **Jordi García-Fernández:** Writing & Review, Evolutionary focus of the article.

## References

1. Afelt, A., Devaux, C., Serra-Cobo, J. & Frutos, R. Bats, bat-borne viruses, and environmental changes. In Bats (ed. Mikkola, H.) 10.5772/intechopen.74377 (IntechOpen, 2018).

2. Aicher, S. M., et al. Species-specific molecular barriers to SARS-CoV-2 replication in bat cells. J. Virol. 10.1128/jvi.00608-22 (2022).

3. Altschul, S. F., Gish, W., Miller, W., Myers, E. W. & Lipman, D. J. Basic local alignment search tool. J. Mol. Biol. 215, 403–410 (1990). 10.1016/S0022-2836(05)80360-2

4. Anthony, S. J. et al. Coronaviruses in bats from Mexico. J. Gen. Virol. 94, 1028–1038 (2013). 10.1099/vir.0.049759-0

5. Ar Gouilh, M., et al. SARS-CoV related betacoronavirus and diverse alphacoronavirus members found in western old-world. Virology 517, 88–97 (2018). 10.1016/j.virol.2018.01.014

6. Beaudette, F. R. & Hudson, C. B. Cultivation of the virus of infectious bronchitis. J. Am. Vet. Med. Assoc. 90, 51–58 (1937).

7. Carcereny, A., et al. Detection, quantification, and typing of enteroviruses in wastewater. Protoc. Exch. 10.21203/rs.3.pex-2622/v1 (2024).

8. Chen, J. et al. Bat-infecting merbecovirus HKU5-CoV lineage 2 can use human ACE2 as a cell entry receptor. Cell 188, 1729–1742 (2025). 10.1016/j.cell.2025.01.042

9. Chen, B., Tian, E. K., He, B. et al. Overview of lethal human coronaviruses. Signal Transduct. Target. Ther. 5, 89 (2020). 10.1038/s41392-020-0190-2

10. Colombi, D., Serra-Cobo, J., Métras, R., Apolloni, A., Poletto, C., López-Roig, M., Bourhy, H. & Colizza, V. Mechanisms for lyssavirus persistence in non-synanthropic bats in Europe: insights from a modeling study. Sci. Rep. 9, 537 (2019). 10.1038/s41598-018-36485-y

11. Corman, V. M. et al. Highly diversified coronaviruses in neotropical bats. J. Gen. Virol. 94, 1984–1994 (2013). 10.1099/vir.0.054841-0

12. Cui, J. et al. Evolutionary relationships between bat coronaviruses and their hosts. Emerg. Infect. Dis. 13, 1526–1532 (2007). 10.3201/eid1310.070448

13. Drexler, J. F. et al. Genomic characterization of severe acute respiratory syndrome-related coronavirus in European bats and classification of coronaviruses based on partial RNA-dependent RNA polymerase gene sequences. J. Virol. 84, 11336– 11349 (2010). 10.1128/jvi.00650-10

14. Drexler, J. F., Corman, V. M. & Drosten, C. Ecology, evolution and classification of bat coronaviruses in the aftermath of SARS. Antiviral Res. 101, 45–56 (2014). 10.1016/j.antiviral.2013.10.013

15. Graham, R. L. & Baric, R. S. Recombination, reservoirs, and the modular spike: mechanisms of coronavirus cross-species transmission. J. Virol. 84, 3134–3146 (2010). 10.1128/JVI.01394-09

16. Hall, T. BioEdit: a user-friendly biological sequence alignment editor and analysis program for Windows 95/98/NT. 10.14601/PHYTOPATHOL_MEDITERR-14998U1.29 (1999).

17. Han, Y. et al. Panoramic analysis of coronaviruses carried by representative bat species in Southern China to better understand the coronavirus sphere. Nat. Commun. 14, 5537 (2023). 10.1038/s41467-023-41264-z

18. Hayman, D. T. S., Fooks, A. R., Marston, D. A. & Garcia-R, J. C. The global phylogeography of lyssaviruses-challenging the ‘Out of Africa’ hypothesis. PLoS Negl. Trop. Dis. 10, e0005266 (2016). 10.1371/journal.pntd.0005266

19. Hutterer, R., Ivanova, T., Meyer-Cords, C. & Rodrigues, L. Bat Migrations in Europe: A Review of Banding Data and Literature (BfN-Schriftenvertrieb im Landwirtschaftsverlag, Münster, Germany, 2005).

20. Kesheh, M. M., Hosseini, P., Soltani, S. & Zandi, M. An overview on the seven pathogenic human coronaviruses. Rev. Med. Virol. 32, e2282 (2022). 10.1002/rmv.2282

21. Katoh, K., Misawa, K., Kuma, K. & Miyata, T. MAFFT: a novel method for rapid multiple sequence alignment based on fast Fourier transform. Nucleic Acids Res. 30, 3059–3066 (2002). 10.1093/NAR/GKF436

22. Knipe, D. M. & Howley, P. M. Fields Virology. 6th edition (Lippincott Williams & Wilkins, 2013).

23. Latinne, A. et al. Origin and cross-species transmission of bat coronaviruses in China. Nat. Commun. (2024). 10.1038/s41467-024-55384-7

24. Leopardi, S., Holmes, E. C., Gastaldelli, M., Tassoni, L., Priori, P., Scaravelli, D., Zamperin, G. & De Benedictis, P. Interplay between co-divergence and cross-species transmission in the evolutionary history of bat coronaviruses. Infect. Genet. Evol. 58, 279–289 (2018).

25. Moraga-Fernández, A. et al. A study of viral pathogens in bat species in the Iberian Peninsula: identification of new coronavirus genetic variants. Int. J. Vet. Sci. Med. 10, 100–110 (2022). 10.1080/23144599.2022.2139985

26. Morales, A. E. et al. Bat genomes illuminate adaptations to viral tolerance and disease resistance. Nature 638, 449–458 (2025). 10.1038/s41586-024-08471-0

27. Pons-Salort, M., Serra-Cobo, J., Jay, F., López-Roig, M., Lavenir, R., Guillemot, D., Letort, V., Bourhy, H. & Opatowski, L. Insights into persistence mechanisms of a zoonotic virus in bat colonies using a multispecies metapopulation model. PLoS ONE 9, e95610 (2014). 10.1371/journal.pone.0095610

28. Posada, D. & Crandall, K. A. MODELTEST: testing the model of DNA substitution. Bioinformatics 14, 817–818 (1998). 10.1093/BIOINFORMATICS/14.9.817

29. Puechmaille, S. J. et al. The evolution of sensory divergence in the context of limited gene flow in the bumblebee bat. Nat. Commun. 2, 573 (2011). 10.1038/ncomms1582

30. Serra-Cobo, J., Sanz-Trullén, V. & Martinez-Rica, J. P. Migratory movements of Miniopterus schreibersii in the north-east of Spain. Acta Theriol. 43, 271–283 (1998).

31. Serra-Cobo, J. & López-Roig, M. Bats and emerging infections: an ecological and virological puzzle. Adv. Microbiol. Infect. Dis. Public Health 972, 35–48 (2016). 10.1007/5584_2016_131

32. Shannon, P. et al. Cytoscape: a software environment for integrated models of biomolecular interaction networks. Genome Res. 13, 2498–2504 (2003). 10.1101/gr.1239303

33. Smith, C. S. et al. Coronavirus infection and diversity in bats in the Australasian region. EcoHealth 13, 72–82 (2016). 10.1007/s10393-016-1116-x

34. Soriano-Tordera, C., Buigues, J., Viñals, A., Muscolino, E., Martínez-Recio, R., Díez, J., Monrós, J., Cuevas, J. M., Dufloo, J. & Sanjuán, R. Identification and characterization of novel bat coronaviruses in Spain. PLoS Pathog. 21, e1013371 (2025). 10.1371/journal.ppat.1013371

35. Tamura, K., Stecher, G. & Kumar, S. MEGA11: molecular evolutionary genetics analysis version 11. Mol. Biol. Evol. 38, 3022–3027 (2021). 10.1093/molbev/msab120

36. Tang, X. C. et al. Prevalence and genetic diversity of coronaviruses in bats from China. J. Virol. 80, 7481–7490 (2006). 10.1128/jvi.00697-06

37. Trifinopoulos, J., Nguyen, L.-T., von Haeseler, A. & Minh, B. Q. W-IQ-TREE: a fast online phylogenetic tool for maximum likelihood analysis. Nucleic Acids Res. 44, W232–W235 (2016). 10.1093/nar/gkw256

38. Waterhouse, A. M., Procter, J. B., Martin, D. M. A., Clamp, M. & Barton, G. J. Jalview Version 2-a multiple sequence alignment editor and analysis workbench. Bioinformatics 25, 1189–1191 (2009). 10.1093/bioinformatics/btp033

39. Wong, A. C. P., Li, X., Lau, S. K. P. & Woo, P. C. Y. Global epidemiology of bat coronaviruses. Viruses 11, 174 (2019). 10.3390/v11020174

40. Zamperin, G., Festa, F., Palumbo, E., Quaranta, E., Monne, I., Terregino, C., De Benedictis, P. & Leopardi, S. Discovery of a coronavirus in the Eurasian badger (Meles meles) belonging to a putative new genus. Infect. Genet. Evol. 10.1016/j.meegid.2023.105406 (2023).

41. Zhang, C., Gu, H., Peng, J., Be, H., Liu, Y., Yan, X., Feng, J. & Liu, Y. Phylogenetic relationships and species composition of host community influence the transmission of coronaviruses in sympatric bats. Mol. Phylogenet. Evol. 207 (2025). 10.1016/j.ympev.2025.108343

42. Zhou, P., et al. Contraction of the type I IFN locus and unusual constitutive expression of IFN-α in bats. Proc. Natl Acad. Sci. USA 113(10), 2696–2701 (2016). 10.1073/pnas.1518240113

43. Zhou, S., Liu, B., Han, Y., Wang, Y., Chen, L., Wu, Z. & Yang, J. ZOVER: the database of zoonotic and vector-borne viruses. Nucleic Acids Res. 50, 943–949 (2022). 10.1093/nar/gkab862

